## Supplemental Tables for "The thrombopoietin receptor agonist eltrombopag inhibits human cytomegalovirus replication via iron chelation"

**Suppl. Table 1**. Effects of eltrombopag addition at different time points during the HCMV replication cycle using human foreskin fibroblasts infected with HCMV strain Hi91 at an MOI of 0.02.

| **Time of drug addition** | **$IC_{50}$[1] (nM)** |
|---|---|
| Continuous starting from virus infection | 415 ± 197 |
| 24h pre-treatment | > 10 |
| During the 1h virus adsorption period | 8,979 ± 1153 |
| 1h post infection | 233 ± 35 |
| 24h post infection | 358 ± 105 |
| 48h post infection | 2,493 ± 795 |

[1] Concentration that reduces HCMV late antigen expression by 50%

**Suppl. Table 2**. Effects of eltrombopag on HCMV late antigen (LA) expression in different cell types infected with different virus strains and isolates as determined 120h post infection. Concentrations that reduce LA expression by 50% ($IC_{50}$) are provided. The investigated eltrombopag concentrations did not affect cell viability.

| | **Eltrombopag $IC_{50}$ (nM)** | |
|---|---|---|
| **Virus strain** | **HFFs** | **ASCs** |
| Hi91 | 415 ± 197 | 4331 ± 1450 |
| Davis | 1182 ± 38 | 265 ± 77 |
| Towne | 4230 ± 257 | 2303 ± 616 |
| U1 | 110 ± 7 | 99 ± 30 |
| U59 | 3372 ± 1253 | 296 ± 84 |
| U75 | 389 ± 93 | 1828 ± 491 |
